# The vulnerability of working memory to distraction is rhythmic

**DOI:** 10.1101/2020.02.26.965848

**Authors:** Malte Wöstmann, Troby Ka-Yan Lui, Kai-Hendrik Friese, Jens Kreitewolf, Malte Naujokat, Jonas Obleser

## Abstract

Recent research posits that the cognitive system samples target stimuli in a rhythmic fashion, characterized by target detection fluctuating at frequencies of ~3–8 Hz. Besides prioritized encoding of targets, a key cognitive function is the protection of working memory from distractor intrusion. Here, we test to which degree the vulnerability of working memory to distraction is rhythmic. In an Irrelevant-Speech Task, *N* = 23 human participants had to retain the serial order of nine numbers in working memory while being distracted by task-irrelevant speech with variable temporal onsets. The magnitude of the distractor-evoked N1 component in the event-related potential as well as behavioural recall accuracy, both measures of memory distraction, were periodically modulated by distractor onset time in approximately 2–4 cycles per second (Hz). Critically, an underlying 2.5-Hz rhythm explained variation in both measures of distraction such that stronger phasic distractor encoding mediated lower phasic memory recall accuracy. In a behavioural follow-up experiment, we tested whether these results would replicate in a task design without rhythmic presentation of target items. Participants (*N* = 6 with on average >2,500 trials, each) retained two line-figures in memory while being distracted by acoustic noise of varying onset across trials. In agreement with the main experiment, the temporal onset of the distractor periodically modulated memory performance. These results suggest that during working memory retention, the human cognitive system implements distractor suppression in a temporally dynamic fashion, reflected in ~400-ms long cycles of high versus low distractibility.

## Introduction

Since our environment is rich in sensory stimuli, our attentional system faces two closely intertwined, yet conceptually dissociable tasks: First, relevant target stimuli need to be prioritized by attention in order to encode them into working memory. Second, irrelevant distractors need to be suppressed to avoid working memory interference. The cognitive system has to orchestrate these two processes dynamically in time (for review, see Nobre & van Ede, 2018), in order to prioritize targets and to suppress distractors from the ongoing stream of sensory inputs. While recent research suggests that the attentional sampling of target stimuli is not constant throughout time but periodically modulated at frequencies in the range of 3–8 Hz (for review, see Fiebelkorn & Kastner, 2019), it is at present unclear whether the vulnerability to distraction is rhythmic as well.

It is long known that rhythmic sensory stimulation has the potency to entrain so-called “attending rhythms” (Large & Jones, 1999), which eventually results in enhanced target stimulus detection at time points in-phase versus out-of-phase with the entrained rhythm (in vision: de Graaf et al., 2013; in audition: M. R. Jones, Johnston, & Puente, 2006). Studies have also revealed the neural basis of this phenomenon: Entrainment of brain oscillations by rhythmic stimulation relates to phasic modulation of target detection (Henry & Obleser, 2012; Spaak, de Lange, & Jensen, 2014). Furthermore, there is some evidence that direct entrainment of neural oscillations by transcranial alternating current stimulation (tACS) induces phasic modulation of target stimulus detection (Gundlach, Muller, Nierhaus, Villringer, & Sehm, 2016; Helfrich et al., 2014). An important interim summary of this research is that the sensory gain for processing to-be-attended target stimuli fluctuates periodically, likely so as a result of neural entrainment (for recent review articles, see Lakatos, Gross, & Thut, 2019; Obleser & Kayser, 2019).

Interestingly, several studies have demonstrated that target detection fluctuates rhythmically even in the absence of entraining sensory input or rhythmic brain stimulation (e.g. Fiebelkorn, Saalmann, & Kastner, 2013; Landau & Fries, 2012). In essence, these studies found that the time interval between an alerting stimulus and subsequent target presentation periodically modulated target detection at frequencies in the theta range of 3–8 Hz. On the neural level, rhythmic attentional sampling was accompanied by theta-rhythmic modulation of gamma responses (Landau, Schreyer, van Pelt, & Fries, 2015) and frontoparietal theta activity in humans (Helfrich et al., 2018) and non-human primates (Fiebelkorn, Pinsk, & Kastner, 2018). These results suggest that attention samples target stimuli at a theta rhythm.

Attention and working memory are tightly coupled neuro-cognitive faculties (Awh, Vogel, & Oh, 2006). This is evidenced also by retro-cueing paradigms that require the flexible allocation of attention to items in memory (e.g. Lim, Wöstmann, & Obleser, 2015; Oberauer & Hein, 2012; Schneider, Mertes, & Wascher, 2016). Furthermore, cued attention – which is by now the most prevalent paradigm to reveal rhythmic sampling – can be considered closely related to working memory, in agreement with the view that attention and working memory pose a unitary system (Postle, 2006). It is thus straight-forward to assume that sampling of target objects in memory might be theta-rhythmic as well (Fiebelkorn & Kastner, 2019). Speaking directly to this, Peters and colleagues (2018) varied the time-interval between a retro-cue (which cued one of two spatially separated visual objects in memory) and a subsequent memory probe. They found rhythmic modulation of response times to the probe at a frequency of 6 Hz. Interestingly, rhythmic trajectories of response times across probe times for the cued versus non-cued object were in anti-phase, suggesting that target items in working memory were sampled in alternation. From a more general perspective, theta oscillations have been implicated in working memory function as they increase during temporal order maintenance (Hsieh, Ekstrom, & Ranganath, 2011) and induce increased working memory performance (for a recent review, see Hanslmayr, Axmacher, & Inman, 2019) if entrained by rhythmic transcranial magnetic stimulation (Albouy, Weiss, Baillet, & Zatorre, 2017) or by rhythmic visual stimulation (Köster, Martens, & Gruber, 2019).

Here, we test whether the temporal onset of task-irrelevant auditory stimuli would prove disruptive in a slow oscillatory fashion. During working memory retention, a single auditory distractor stimulus was presented at varying onset times across trials. Distractor onset periodically modulated both, the distractor-evoked N1 component in the event-related potential (at ~3.5 Hz) and memory recall accuracy (at ~2 Hz). Critically, these two measures of distraction were co-modulated by an underlying ~2.5-Hz rhythm. In a behavioural follow-up experiment, we conceptually replicated the rhythmic vulnerability of working memory to distraction in a visual match-to-sample paradigm.

## Materials and methods

The main study poses a re-analysis of a previously published dataset (Wöstmann, Lim, & Obleser, 2017). Below, we describe essential methodological aspects but we refer to the original article for details.

### Participants

In the main study and in the follow-up experiment, we tested *N* = 23 (12 females & 11 males; mean age: 24.5 years) and *N* = 6 (5 females & 1 male; mean age: 22 years) participants, respectively. Participants provided informed consent and were either compensated financially or received course credit. Experimental procedures were approved by the ethics committee of the University of Lübeck.

### Task design and stimuli in the main study

For the main study, to-be-remembered auditory materials were the German numbers from 1 to 9, spoken by a female voice (average number duration 0.6 s). To-be-ignored distractors were short German sentences (5–8 words, average duration: 2.1 s), taken from a German version of speech-in-noise sentences (Erb, Henry, Eisner, & Obleser, 2012), spoken by the same female voice as the numbers.

We used an adapted version of the Irrelevant-Speech Task (Colle & Welsh, 1976; Jones & Morris, 1992). On each trial, participants had to retain nine target numbers in memory, which were presented in random order with onset-to-onset delay of 0.75 s (presentation rate of 1.33 Hz). The most important manipulation for the present study was the temporal onset of the speech distractor. During the ensuing 5-s memory retention period, a task-irrelevant distractor sentence was presented, which had to be ignored. On each trial, the distractor onset was drawn from a uniform distribution including all millisecond values between 1.035 and 1.835 s relative to the offset of the last target number. On average, the distractor sentence was centred in the middle of the retention period, which corresponded to a distractor onset of 1.435 s.

At the end of each trial, participants were presented with a number pad on a computer screen. They had to select the numbers in their order of presentation using a computer mouse. In order to prevent participants’ motor preparation for a particular behavioural response, the number pad was randomly arranged on each trial.

The experiment was implemented in Presentation software (Neurobehavioral Systems). All auditory materials were presented via Sennheiser HD-25 headphones at a comfortable level of ~65 dB A.

### Additional manipulations of no interest in the main study

Two additional manipulations of the speech distractor were implemented in the main study but these have been analysed previously (Wöstmann et al., 2017) and were thus not of primary interest here. First, acoustic detail of distractor sentences was manipulated using noise-vocoding. In brief, noise-vocoding parametrically degrades the spectral content of the speech signal but leaves the temporal information largely intact (Shannon, Zeng, Kamath, Wygonski, & Ekelid, 1995). The fewer spectral bands are used for noise-vocoding, the stronger the degradation and the lower the resulting intelligibility of speech materials. In the present study, 1, 4, and 32 spectral bands were used for vocoding. As we know from previous studies that a logarithmic increase in spectral bands of noise-vocoded distractors linearly increases the degree of distraction (Wöstmann et al., 2017; Wöstmann & Obleser, 2016), we regressed out noise-vocoding (coded 1, 2, and 3 for 1, 4, and 32 spectral bands, respectively) in the statistical analysis (see below).

Second, half of the distractor sentences ended in a final word that was highly predictable (e.g. “She covers the bed with fresh sheets”; translated from German), whereas the final word was unpredictable in the other half (e.g. “We are very happy about the sheets”). Since we found in a previous study (Wöstmann & Obleser, 2016) and in the current dataset (Wöstmann et al., 2017) that final word predictability of distractor sentences had no effect on performance in the Irrelevant-Speech Task, predictability was not further considered in the present study.

### EEG recording and preprocessing in the main study

The EEG was recorded at 64 active scalp electrodes (Ag/Ag-Cl; ActiChamp, Brain Products, München, Germany) at a sampling rate of 1,000 Hz, with a DC–280 Hz bandwidth, against a left-mastoid reference (channel TP9). All electrode impedances were kept below 5 kOhm. Offline, the continuous EEG data were filtered (0.3-Hz high-pass; 180-Hz low-pass) and segmented into epochs relative to the onset of the first target number (–2 to +16 s). As described in detail in (Wöstmann et al., 2017), an independent component analysis (ICA) and subsequent automatic artefact rejection were used to clean the EEG data.

For the purpose of the present study, the single-trial epoched EEG data were aligned to the onset of the task-irrelevant speech distractor (–2 to +4 s), re-referenced to the average of all electrodes, low-pass filtered at 10 Hz in order to obtain robust single-trial distractor-evoked ERPs, and baseline corrected by subtraction of mean voltage in the time window –0.1 to 0 s relative to distractor onset. For the EEG data analyses we used the Fieldtrip toolbox (version 2018-06-14; Oostenveld, Fries, Maris, & Schoffelen, 2011) for Matlab (R2018a) and customized Matlab scripts.

### Analysis of periodic modulation of recall behaviour and distractor-evoked N1 amplitude in the main study

Each participant performed 180 experimental trials. After removal of artefactual EEG epochs, an average of 153 trials (SD = 23.4) per participant remained for the analyses of behavioural and EEG data.

The two major outcome variables of interest were memory recall accuracy and neural distractor encoding. The former was quantified as the proportion of numbers recalled at their respective positions of presentation; the latter was quantified as the mean EEG amplitude at electrode Cz in the time interval 0.09–0.13 s relative to distractor onset. This time interval was determined by inspection of the grand-average ERP waveform (see Fig. 1D). To quantify and visualize memory recall and N1 amplitude as a function of distractor onset time, both of these single-trial measures were averaged for a 0.1-s long sliding window, which was moved in 0.01-s steps over distractor onset time for each participant, resulting in 71 time bins (see Fig. 1D). For single participants, each of these overlapping time bins included on average 19.13 trials (SD: 0.71).

**Figure 1.**
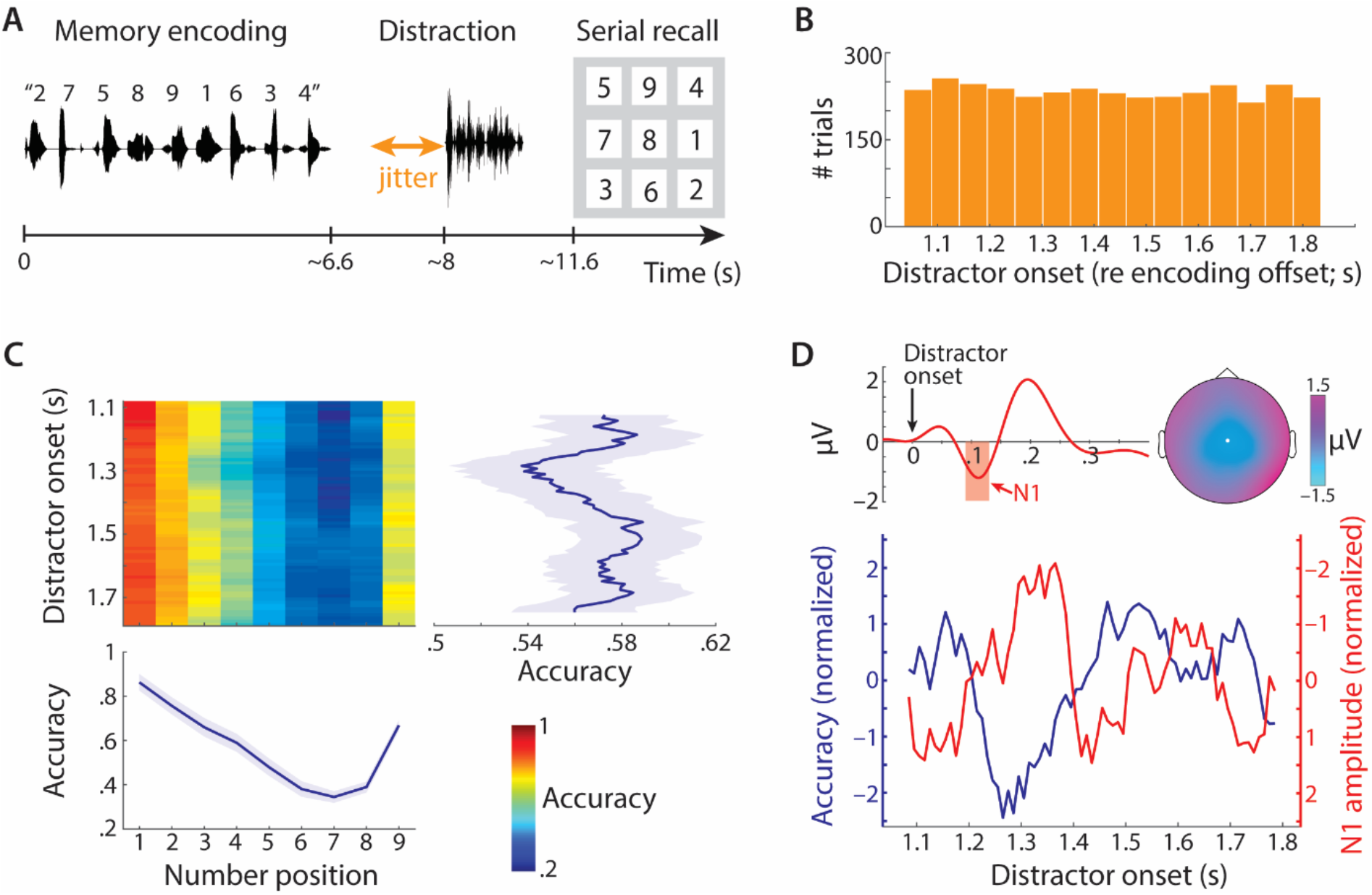
(**A**) Irrelevant-Speech Task. During retention of numbers in memory, participants were distracted by a task-irrelevant spoken sentence. At the end of each trial, participants had the task to select the numbers from a visual display in the order of presentation. (**B**) Histogram of distractor onset times across all (N = 23) participants. (**C**) Heatmap shows average recall accuracy as a function of number position (x-axis) and binned distractor onset time (y-axis). Blue lines show marginal means. Shaded areas show ±1 between-subject standard error of the mean (SEM). (**D**) Top: Grand-average distractor-evoked event-related potential (ERP) at electrode Cz. The topographic map shows the N1 component in the time window .09–.13 s after distractor onset (red shaded area). Bottom: Lines show z-transformed grand-average proportion correct (blue) and N1 amplitude (red) as a function of binned distractor onset time. Note that negative N1 amplitude values (referring to stronger distractor encoding) are plotted upwards.

To quantify rhythmic modulation of memory recall and distractor-evoked N1 across distractor onset time, linear mixed-effects models were used to regress single-trial outcome measures on circular sine and cosine predictors (a method that also proved superior in a recent article that compared different approaches to analyse phasic modulation of neural and behavioural responses; Zoefel, Davis, Valente, & Riecke, 2019).

Before the analysis of rhythmic memory recall modulation, single-trial accuracy (ranging from 0 to 9 out of 9 correctly recalled numbers) was logit transformed. Since the logit for proportions 0 and 1 is undefined, the following adjusted equation was used to logit-transform single-trial proportion correct (PC) values (Fox & Weisberg, 2019):

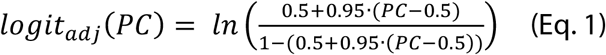

Prior to mixed-effects model analyses, predictor and outcome variables were z-transformed.

We first regressed out potential effects of auditory distractor vocoding, by regression of single-trial accuracy and N1 amplitude on vocoding level plus random subject intercept, using the *fitlme* function in Matlab and the model formulae ‘Accuracy ~ Vocoding + (1|Subject ID)’ and ‘N1 amplitude ~ Vocoding + (1|Subject ID)’, respectively. Next, we regressed the residuals of these regression models on sine- and cosine transformed distractor onset times, using the model formulae ‘Accuracy residuals ~ Sine predictor + Cosine predictor + (1ļSubject ID)’ and ‘N1 residuals ~ Sine predictor + Cosine predictor + (1ļSubject ID)’. Sine- and cosine-transformed distractor onsets for a given frequency *freq* were determined using the formulae:

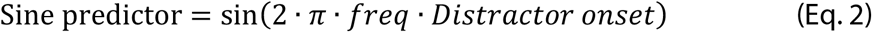

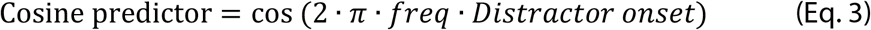

This step was iteratively repeated for frequencies 0.5 to 8 Hz in steps of 0.25 Hz. For each frequency, spectral magnitude was computed from the resulting coefficients (coef) for the Sine and Cosine predictor using the formula:

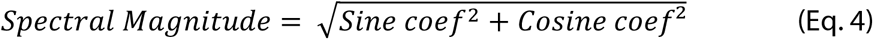

Significance of rhythmic modulation of measures of distraction (accuracy & N1 amplitude) was derived using a permutation approach. For each frequency, we compared the empirical value of spectral magnitude to 5,000 permutations of the data, generated by randomly shuffling single-trial sine- and cosine-transformed distractor onset time within individual participants. Spectral magnitude was considered significantly larger than expected under the null hypothesis if the empirical value of spectral magnitude for a given frequency exceeded the 95^th^ percentile of permuted data (i.e. one-sided testing).

### Analysis of joint modulation of behaviour and N1 amplitude in the main study

We computed the cross-correlation (using the *xcorr* function in Matlab) of average binned accuracy and N1 amplitude as a function of distractor onset time. We also calculated the cross power spectral density of these two measures (using the *cpsd* function in Matlab; rectangular window length = 0.3 s, window overlap = 0.1 s). Empirical cross-correlation and cross power spectral density were compared to surrogate data, generated by 5,000 permutations of single-trial distractor onset time within single participants before binning the data.

In order to test whether a joint 2.5-Hz rhythm, which turned out most prominent in the cross power spectral density analysis, modulates memory recall accuracy directly or indirectly (mediated via N1 amplitude), we performed a mediation analysis. To this end, we extracted the group-level 2.5-Hz phase (*φ_2.5Hz_*) of the mixed-effects model to regress single trial accuracy on sine-and cosine-transformed distractor onset time (using the *atan* function in Matlab to calculate the inverse tangent) using the formula:

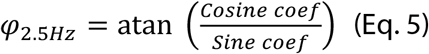

Three group-level, time-domain signals were entered into the mediation analysis: average accuracy binned for distractor onset time, average N1 amplitude binned for distractor onset time, and the 2.5-Hz rhythm with phase *φ_2.5 Hz_*. To estimate individual path coefficients between the 2.5-Hz rhythm, average accuracy, and N1 amplitude, linear regression analyses were used (using the *fitlm* function in Matlab). Significance of the *indirect path*, which would link the 2.5-Hz rhythm and memory recall accuracy via the mediator distractor-evoked N1 amplitude, was calculated in the *medmod* package in Jamovi (version 1.1.9).

### Task design, stimuli and statistical analysis in the follow-up experiment

In order to test whether the results of the main experiment would conceptually replicate in a task that does not contain rhythmic presentation of to-be-remembered target stimuli, we used a visual match-to-sample task in a behavioural follow-up experiment. On each trial, participants were presented with a central black fixation cross on a screen with grey background, flanked by two line-figures, each one of which was generated by connecting 20 randomly drawn points with black lines. Edge length of line-figures was ~4.05 * 4.05 cm and each line figure was displaced from the central fixation cross by ~0.405 cm. In order that the entire visual display did not exceed 5° visual angle, participants were seated at a distance of ~102 cm from the screen.

Participants had the task to encode both line-figures into visual working memory while remaining central fixation. Line-figures were presented for a duration of 0.3 s. In order to reset the phase of a potential rhythm that samples the two line-figures in memory, the corners of one of the two line-figures were highlighted in white within the last 0.05 s of the encoding period. Note that such a “flash event” was also used in previous studies that tested the rhythmic attentional sampling of visual target objects (e.g., Fiebelkorn et al., 2013; Landau & Fries, 2012).

During the ensuing 3-s retention period, participants kept the line-figures in memory while being distracted by a broad-band noise distractor (0.1–5 kHz) of 1-s duration, with 0.01-s linear onset and offset ramps. The distractor onset varied across 24 linearly spaced time points in the time interval 0.5–1.5 s relative to the end of the encoding period. In the end of a trial, one line-figure was presented on the left or right side of the fixation cross. Participants had to indicate via button press whether the line-figure matched the one presented on the same side during encoding. Note that the follow-up experiment also contained trials without a distractor stimulus, as well as trials with the flash event on the side opposite to the probed line-figure. These trials were excluded from the present analysis. The experiment was implemented in Psychtoolbox (Brainard, 1997) for Matlab. Auditory materials were presented via Sennheiser HD-280 Pro headphones.

The follow-up experiment was designed to increase the signal-to-noise ratio in the data by testing relatively few participants on a huge number of experimental trials. Participants *(N* = 6) were tested in two data recording sessions of ~3.5–4 h duration, each (on two different days). For one participant, data of the first session were excluded due to a misunderstanding of task instructions. One other participant did not finish the experiment, so that 77 trials were missing. On average, 2,537 trials (SD: 566) of each participant were included in the present analysis, corresponding to an average of 105.7 trials per participant and distractor onset time. Note that the approach to collect rich behavioural data of few individuals has been adopted also in previous studies that investigated rhythmic sampling of target stimuli in monkeys (Fiebelkorn et al., 2018) and in humans (Tomassini, Spinelli, Jacono, Sandini, & Morrone, 2015).

For the statistical analysis, we used the same procedure as for the main study with the following exceptions. First, no factors were regressed out prior to regression of accuracy on sine- and cosine-transformed distractor onset time. Second, since single-trial accuracy was binary (1 = correct response; 0 = incorrect response), we used logistic regression (using the *fitglme* function in Matlab for a *binomial* distribution of the response variable and a *logit* link function).

For visualization purposes, average accuracy was calculated for each participant and distractor onset time, followed by smoothing via replacement of accuracy for bin_*b*_ by: 0.25 * accuracy bin_*b*−1_ + 0.25 * accuracy bin_*b*+1_ + 0.5 * accuracy bin_*b*_. Then, smoothed accuracy data were averaged across participants, detrended, and z-transformed.

### Circular statistics

In order to derive single-subject phase estimates of periodic modulation of accuracy and N1 amplitude (Fig. 3A and Fig. 5), we performed Fast Fourier Transforms (rectangular window; zero-padding to derive a frequency resolution of 0.05 Hz) on single-subject time courses of accuracy and N1 amplitude binned for distractor onset time. To calculate the circular mean, phase lag, and resultant vector length, we used the respective functions *circ_mean, circ_dist*, and *circ_r*, as implemented in the circular statistics toolbox for Matlab (Berens, 2009).

**Figure 2.**
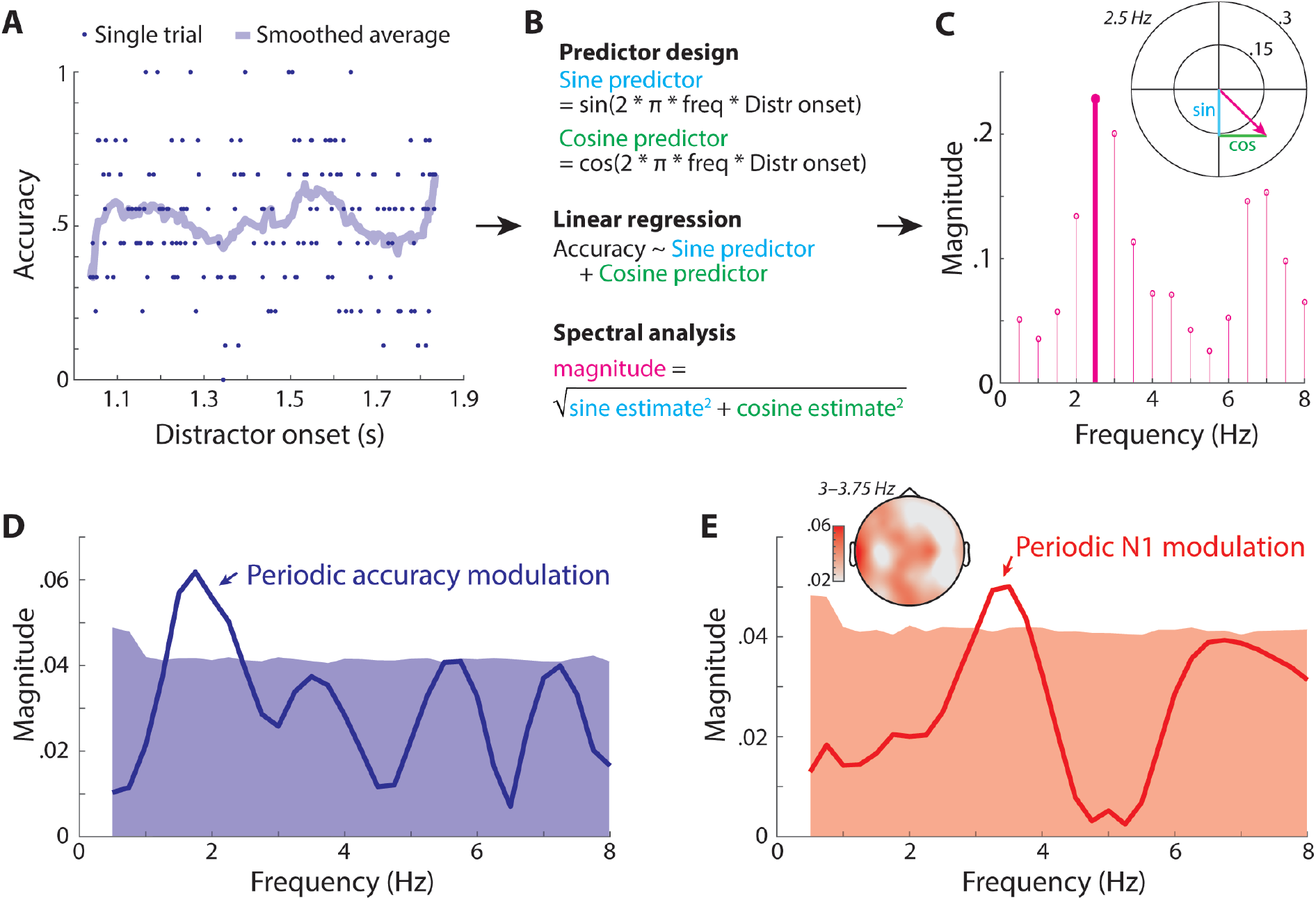
(**A–C**) Schematic depiction of the analysis procedure for periodic modulation of single-trial measures of distraction. (**A**) Dots show single-trial accuracy as function of distractor onset for one exemplary participant. For visualization, the shaded line shows accuracy averaged across twenty neighbouring distractor onsets. (**B**) For statistical analysis, sine- and cosine-transforms of distractor onset time were used as predictors in a linear regression to model single-trial accuracy. Spectral magnitude was calculated as the square root of the sum of squared sine and cosine estimates. Variables with uppercase names denote vectors that contain multiple entries; lowercase variables contain a single entry. (**C**) Spectral magnitudes derived from multiple linear models for frequencies 0.5–8 Hz. The unit circle shows sine/cosine estimates and spectral magnitude (length of pink arrow) for a frequency of 2.5 Hz. (**D**) Solid line shows spectral magnitude of periodic modulation of single-trial accuracy by distractor onset (derived from linear mixed-effects models). Shaded area shows the 95^th^ percentile of surrogate spectra (derived from 5,000 permutations of single-trial distractor onset time within single participants). (**E**) Same as D but for periodic modulation of N1 amplitude. The topographic map shows the spatial distribution of spectral magnitude averaged across frequencies 3–3.75 Hz.

**Figure 3.**
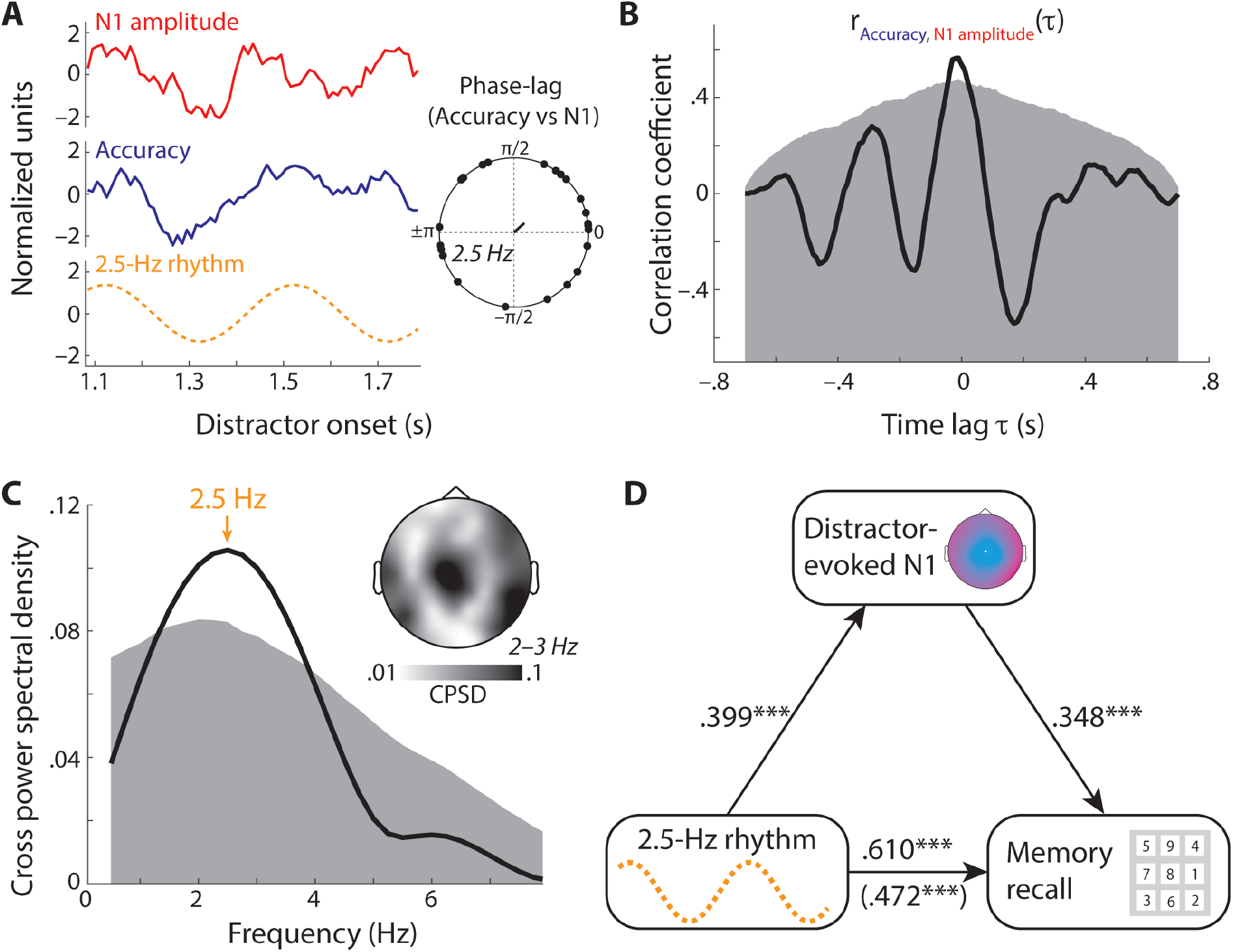
(**A**) Lines show average N1 amplitude (red), accuracy (blue), and the 2.5-Hz rhythm (orange) extracted from the mixed-effects model to regress single-trial accuracy on 2.5-Hz sine- and cosine-transformed distractor onset time. The circular plot shows the 2.5-Hz phase-lag of accuracy and N1 amplitude per subject (dots) and the resultant vector. (**B**) Line shows cross-correlation of average recall accuracy and N1 amplitude binned for distractor onset time. Shaded area shows 95^th^ percentile of surrogate cross-correlations, computed on surrogate data (derived from 5,000 permutations of single-subject distractor onset times). (**C**) Line shows cross power spectral density of average recall accuracy and N1 amplitude binned for distractor onset time. Shaded area shows the 95^th^ percentile of surrogate data (computed for the same permutations as in B). The topographic map shows the spatial distribution of cross power spectral density averaged across frequencies 2–3 Hz. (**D**) Depiction of mediation analysis. Numbers at arrows show beta-coefficients of linear regression models. The total effect of the 2.5-Hz rhythm on memory recall accuracy (*β* = 0.610) was weakened when N1 amplitude was controlled for (*direct effect; β* = 0.472), resulting in a significant partial mediation via N1 amplitude (*indirecteffect; β* = 0.139). *** *p* < 0.001.

**Figure 4.**
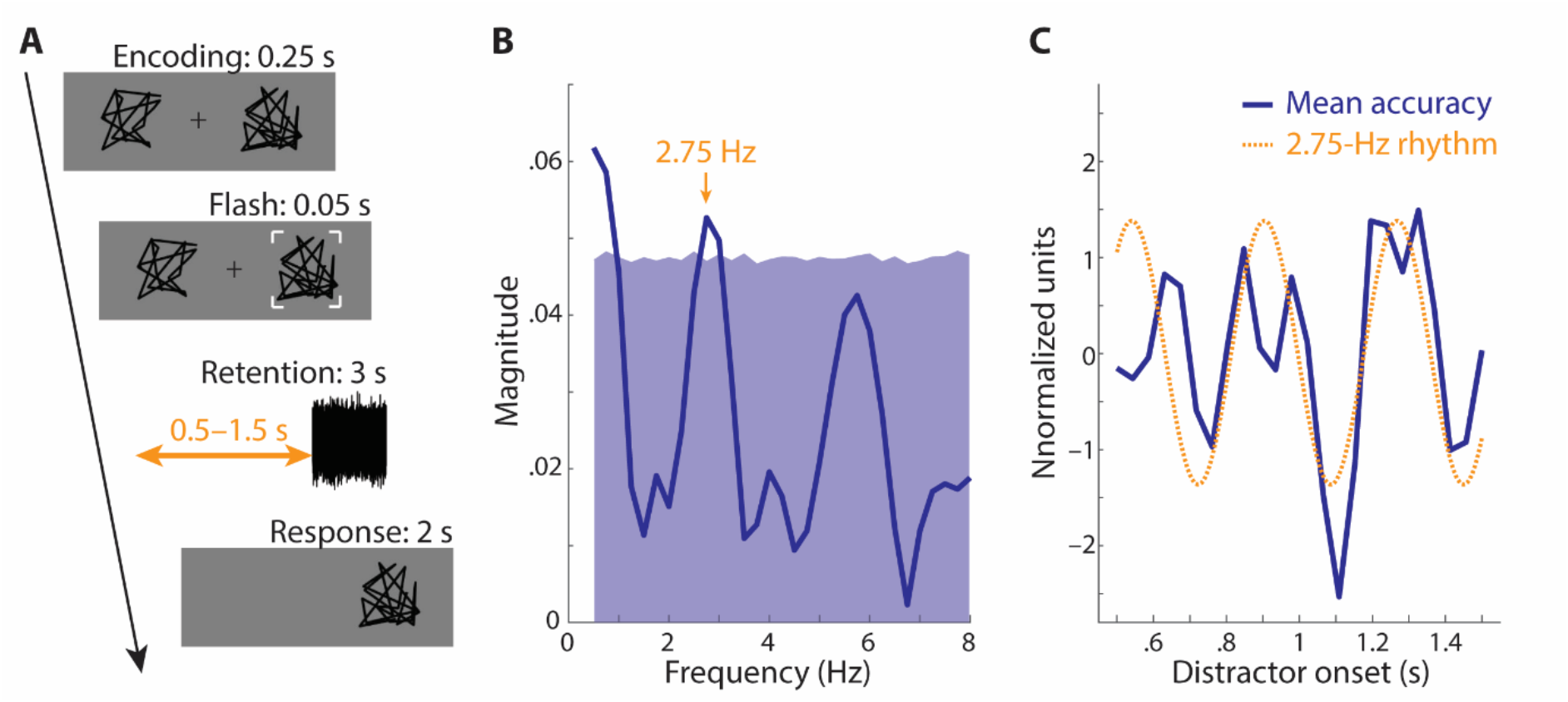
(**A**) Design of behavioural follow-up experiment. Participants (N = 6) encoded two line-figures, before one of them was highlighted by a flash event. During the ensuing 3-s memory retention period, an auditory distractor was presented at one of 24 linearly spaced delays between 0.5 and 1.5 s. In the end of a trial, participants had to indicate whether the probe figure matched the one presented on the same side during encoding (response timeout: 2 s). (**B**) Line shows spectral magnitude of periodic modulation of single-trial accuracy by distractor onset. Shaded area shows the 95^th^ percentile of surrogate spectra (derived from 5,000 permutations of single-trial distractor onset time within single participants). (**C**) The blue line shows average accuracy (which was temporally smoothed, detrended, and z-transformed for purpose of visualization). The orange line shows the 2.75-Hz rhythm derived from the mixed model to regress accuracy on sine- and cosine-transformed distractor onset time.

**Figure 5.**
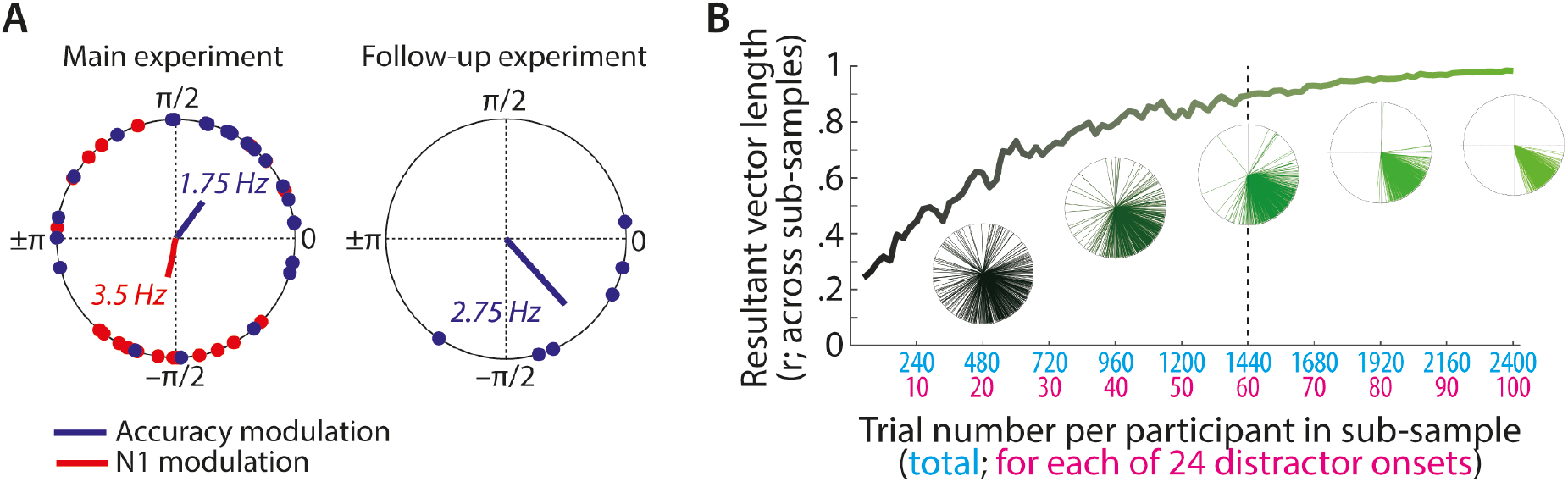
(**A**). Dots show single-subject phase angles for peak frequencies of accuracy modulation (blue; in the main and follow-up experiment) and N1 modulation (red; in the main experiment). Lines show resultant vectors for accuracy modulation in the main experiment (blue; resultant vector length, *r* = 0.383) and follow-up experiment (*r* = 0.735), as well as for N1 modulation in the main experiment (red; *r* = 0.331). (**B**) Circular plots show 500 vectors of unit length. Each vector was derived by sub-sampling the respective number of trials fa-axis) for each participant, computation of the 2.75-Hz phase angle, and calculation of the circular mean across the six participants in the follow-up experiment. Since for one participant only half of the data were used for data analysis (see Materials and Methods for details), sub-samples included up to 48 trials for this participant. The solid line shows the resultant vector length across the 500 sub-samples. Note that the critical information in this figure is at which point no considerable further increase in the resultant vector length (which will always asymptotically reach *r* = 1) can be observed with increasing trial number (dashed line).

## Results and Discussion

In the main study, participants (*N* = 23) retained the serial order of nine numbers in working memory, while being distracted by a task-irrelevant spoken sentence of varying temporal onset across trials (Fig. 1A&B). As it is typical for an Irrelevant-Speech Task (e.g. D. M. Jones, Macken, & Murray, 1993; Kreitewolf, Wöstmann, Tune, Plöchl, & Obleser, 2019; Wöstmann & Obleser, 2016), primacy and recency effects in working memory (Henson, 1998) were reflected in relatively high recall accuracy for early number positions (1–4) and for the last number position, respectively (Fig. 1C).

### Distractor onset periodically modulates target recall and distractor encoding

In order to test whether the vulnerability of working memory to distraction is rhythmic, we investigated two outcome measures of interest. First, on the behavioural level, serial recall accuracy averaged across number positions was thought to be inversely related to the degree of distraction. That is, the more a participant is distracted by irrelevant speech, the lower the recall accuracy on an individual trial. Second, on the neural level, the amplitude of the N1 component of the distractor-evoked ERP was utilized as a measure of neural distractor encoding. The rationale for this was that previous studies using the Irrelevant-Speech Task found that N1 amplitude reflected the degree of distraction (Bell, Dentale, Buchner, & Mayr, 2010; Weisz & Schlittmeier, 2006). N1 amplitude was extracted at electrode Cz in the time window 0.09–0.13 s following distractor onset.

Both average recall accuracy and N1 amplitude fluctuated periodically across distractor onset times (Fig. 1D), with such distractor onsets associated with stronger distractor encoding (more negative N1 amplitude) coinciding with lower recall accuracy. As to be expected for two measures of the same underlying construct (i.e. distraction), average serial recall accuracy and N1 amplitude binned for distractor onset time were significantly correlated (*r* = 0.536; *p* < 0.001).

To statistically analyse the periodic modulation of both measures of distraction, we used linear mixed-effects models to regress single-trial accuracy and N1 amplitude on sine- and cosine-transformed distractor onset time (Fig. 2A–C; see Materials and Methods for details). Empirical spectra from mixed-effects models were contrasted against surrogate spectra, derived from 5,000 permutations of single-subject distractor onset time. We found significant periodic modulation of both measures by distractor onset time, for memory recall accuracy at frequencies ~1.25–2.5 Hz and for N1 amplitude at frequencies ~3–3.75 Hz (Fig. 2D&E).

While previous research has shown that the attentional sampling of exogenous target stimuli (Fiebelkorn et al., 2013; Landau & Fries, 2012) and target stimuli retained in working memory is rhythmic (Peters et al., 2018), we demonstrate here that the vulnerability of working memory to distraction is rhythmic as well. Mechanistically, a straight-forward interpretation of the present results is that the suppression of distractors fluctuates rhythmically, which results in alternating periods of higher versus lower vulnerability to distraction. If the onset of a distractor stimulus falls into a period of high distractibility (i.e. low distractor suppression), neural encoding of the distractor is enhanced, resulting in stronger memory disruption. Such an interpretation implies that distractor encoding and memory recall are co-modulated by a joint rhythm, which we tested next.

### A joint rhythm modulates distractor encoding and memory recall

We tested to what degree memory recall accuracy and the distractor-evoked N1 amplitude were periodically modulated by a joint underlying rhythm. To this end, we employed average accuracy and N1 amplitude binned for distractor onset time (shown in Fig. 1D) and computed their cross-correlation and cross power spectral density, which significantly exceeded surrogate data (derived from 5,000 permutations of single-subject distractor onset time) at frequencies of ~1.5–4 Hz and peaking at 2.5 Hz (Fig. 3). These results are in line with the notion that a common slow-oscillatory rhythm (here, 2.5 Hz) modulates memory recall accuracy and the distractor-evoked N1 amplitude.

If the rhythmic modulation of distractor encoding drives synchronized modulation of memory recall accuracy, the 2.5-Hz periodic modulation of memory recall should be mediated by the distractor-evoked N1 amplitude. To test this, we performed a mediation analysis. We extracted the phase of 2.5-Hz rhythm from the mixed-effects model to regress single-trial accuracy on 2.5-Hz sine- and cosine-transformed distractor onset time (*φ_2.5 Hz_* = 2.79). The 2.5-Hz rhythm significantly predicted N1 amplitude (*β*= 0.399; *p* < 0.001) and memory recall (*total effect; β* = 0.610; *p* < 0.001). Importantly, the latter effect was significantly reduced when the regression model included N1 amplitude as an additional predictor (*direct effect; β* = 0.472; *p* < 0.001). A significant indirect effect (*total effect* – *direct effect; β* = 0.139; *p* = 0.009) revealed that the 2.5-Hz rhythmic modulation of memory recall accuracy was partially mediated by the distractor-evoked N1 amplitude (22.7 % mediation). This supports the view that periodic fluctuation of distractibility induces synchronized states of increased versus decreased neural distractor encoding, which in turn result in increased versus decreased working memory disruption, respectively.

These results provide converging behavioural and neural evidence that a 2.5-Hz rhythm governs the degree to which task-irrelevant stimuli disrupt working memory (see also next section). Notably, however, rhythmic vulnerability to memory distraction found in the present study appears to be somewhat slower than rhythmic sampling of target stimuli, which is typically observed in the theta range of 3–8 Hz (Fiebelkorn & Kastner, 2019). Why would that be?

Recent neuroscience work suggests that distractor suppression is not the mere inverse of target enhancement, but an independent neuro-cognitive mechanism that operates on distractors in the sensory environment (Noonan et al., 2016; Wöstmann, Alavash, & Obleser, 2019), as well as on distractors in working memory (Schneider, Goddertz, Haase, Hickey, & Wascher, 2019). In this view, it is well conceivable that rhythms of different frequency orchestrate the dynamic sampling of target stimuli (at 3–8 Hz) versus suppression of distractors (at ~2.5 Hz).

Alternatively, it might be that the nature of the present experimental paradigm induced rhythmic distractibility at 2.5 Hz. In contrast to previous studies that investigated rhythmic sampling of visual targets, the present study used a serial recall working memory task in the auditory modality. Furthermore, the rhythmic presentation of target stimuli in the present study might have entrained distractibility to a certain extent. To test whether these alternative explanations can be ruled out, we next analysed the data of a behavioural follow-up experiment.

### The origin of rhythmic distractibility

An obvious question is whether the observed rhythmic modulation of memory distractibility is entrained by external sensory stimulation or whether it reflects a spontaneous internal rhythm underlying memory function. Although the present study cannot provide a definite answer to this question, we address both options below.

In theory, it is possible that the rhythmic presentation of target numbers entrained an attending rhythm at 1.33 Hz, which periodically modulated also the distractibility during memory retention. Consistent with this notion, distractor onset time modulated recall accuracy at frequencies including this attending rhythm (~1.25–2.5 Hz). Furthermore, the time point of highest memory distractibility (at ~1.26 s following target offset; see Fig. 1D) falls precisely in-between two peaks of the hypothetical 1.33-Hz attending rhythm (i.e., in-between the onsets of putative number positions 11 and 12). However, the joint periodic modulation of accuracy and distractor-evoked N1 amplitude was strongest at a frequency of 2.5 Hz rather than 1.33 Hz, which speaks against the view of entrained distractibility.

Alternatively, the observed ~2.5-Hz rhythm might be a spontaneous internal rhythm underlying memory function (see Fig 3D). If this is the case, a similar rhythm should be observed also in the absence of a potentially entraining stimulus sequence. In a behavioural follow-up experiment, we used a visual match-to-sample task to test this hypothesis. While participants retained two line-figures in working memory, they were distracted by an auditory noise distractor of varying onset time (Fig. 4). Distractor onset time periodically modulated task accuracy at a frequency of 2.75 Hz, which supports the claim that memory distractibility fluctuates rhythmically even when no rhythmic sensory stimulation precedes working memory retention. Note that in addition to a frequency of 2.75 Hz, accuracy in the follow-up experiment was modulated at lower frequencies (< 1 Hz), most likely because distractor onset time linearly modulated task accuracy. Later distractors were more detrimental to accuracy (linear effect of absolute distractor onset time; *β* = −0.62; *p* = 0.001 in a logistic regression of single-trial accuracy).

The results of this follow-up experiment constitute an analogy of distractor suppression to what has previously been shown for the attentional sampling of target stimuli (Fiebelkorn et al., 2013; Landau & Fries, 2012): there is an inherent (~2.5-Hz) rhythm that regulates the temporally dynamic suppression of task-irrelevant distraction during working memory maintenance. Relatively slow oscillatory effects in the delta frequency band (< 4 Hz) have also been related to the sequential prioritization in working memory (de Vries, van Driel, & Olivers, 2019).

Critically, although the dependent measures of distraction in the present study (i.e., memory recall and N1 amplitude) were both modulated by a salient ~2.5-Hz rhythm, the present study does not disclose the neural origin of this rhythm. Spectral analyses of EEG activity during the memory retention interval did not give any indication of periodic oscillatory activity at a frequency of ~2.5 Hz (data not shown). Therefore, our measures of distraction can be considered instruments to sample the phase of the ~2.5-Hz rhythm. The rhythm as such, however, is not picked up in the EEG in the present study. One candidate neural structure where the inherent rhythm that modulates distractibility might originate is the hippocampus, where endogenous delta oscillations have been found to orchestrate synchronized states of working memory activation (Leszczynski, Fell, & Axmacher, 2015). In this sense, the ~2.5-Hz rhythm of distractor suppression might be a temporally dynamic mechanism specific to working memory. That is, during memory retention of target items, suppression of distraction is not a constant process but instead modulated at ~2.5 Hz, which allows for (partial) memory intrusion of currently task-irrelevant stimuli approximately every 400 ms.

While the present data disclose a periodic modulation of distractor suppression at a frequency of ~2.5-Hz, this does not preclude that this rate of distractor suppression might adapt to task demands. For instance, we found rhythmic modulation of neural indices of attentional filtering during stimulus encoding (i.e. target selection and distractor suppression) in previous studies at 0.67 Hz (Wöstmann, Herrmann, Maess, & Obleser, 2016) and 0.375 Hz (Wöstmann, Schmitt, & Obleser, 2020), corresponding to the respective rates of stimulus presentation.

### Impact of trial number on reliability of periodic memory modulation

The present study includes two experiments, which differ considerably in the number of trials employed to sample the rhythmic modulation of distractor intrusion in working memory. While on average 153 trials per participant were analysed in the main experiment, the follow-up experiment included on average > 2,500 trials per participant.

Since our analysis pipeline was specifically designed to pick up phase-consistent modulation of memory distraction across participants, the degree of between-participant phase-consistency can be considered a measure of reliability. In agreement with the assumption that a larger trial number should decrease the influence of measurement error, phase-consistency of periodic modulation of memory recall accuracy was considerably higher in the follow-up experiment (resultant vector length across single-participant phase angles at 2.75-Hz, *r* = 0.735; Fig. 5A) compared with the main experiment (at 1.75 Hz; *r* = 0.383).

To estimate the impact of increasing trial number on the ability to pick up phase-consistent modulation of memory distractibility, we employed a sub-sampling approach on the data in the follow-up experiment. For increasing trial numbers of 1 to 100 trials for each one of 24 possible distractor onset times, we randomly sub-sampled trials from the existing data, using 500 sub-samples. For each sub-sample and participant, we calculated the 2.75-Hz phase of memory recall accuracy as a function of distractor onset time, followed by calculating the circular mean across participants. The rationale of this analysis is the following: If relatively small trial numbers are sufficient to sample periodic memory distraction robustly, phase-consistency across sub-samples should be high already for small trial numbers. To the contrary, phase-consistency across sub-samples was relatively small (*r* ≤ 0.4) for up to 10 trials for each one of 24 possible distractor onsets (Fig. 5B). Figure 5B suggests that approximately 60 trials per distractor onset time (resulting in 60 trials x 24 distractor onsets = 1,440 trials per participant) would have been sufficient in the present experiment since phase-consistency across sub-samples did not considerably increase further for larger trial numbers.

Although the results of this sub-sampling analysis do not establish a precise lower bound for the number of trials necessary to robustly sample rhythmic memory distraction, they clearly indicate an advantage of designs with larger trial numbers. Future studies, including large trial numbers and larger participant samples, could employ a similar sub-sampling approach to moreover estimate the necessary ratio of trial number to sample size.

## Limitations

We consider three main limitations of the present study. First, the concept of rhythmicity implies that the mechanism of interest, e.g. attentional sampling of targets or vulnerability to distraction, is rhythmically modulated over an extended period of time. However, for practical reasons, the present study varied the distractor onset within relatively short time windows of 0.8 s in the main study and 1 s in the follow-up experiment. In fact, short time windows of investigation are a common limitation also for studies exploring the rhythmic attentional sampling of targets (1.3 s in Fiebelkorn et al., 2018; 0.8 s in Fiebelkorn et al., 2013; 1.2 s in Helfrich et al., 2018; 1 s in Landau & Fries, 2012; 0.8 s in Peters et al., 2018). Thus, although the present results suggest memory distractibility to be a rhythmic process, it is at present an open question to what extent this rhythmicity extends over longer time intervals.

Second, both experiments in the present study employed only a single distractor stimulus per trial during memory retention to sample periodic memory distractibility. An arguably more efficient method would have been to employ temporally extended soundscapes, which have been used to reveal rhythmic perceptual sampling of auditory scenes at frequencies 1–2 Hz by means of reverse correlation (Kayser, 2019). Furthermore, such an approach would allow to study the accumulation of evidence in the distractor stimulus (see Wyart, de Gardelle, Scholl, & Summerfield, 2012).

Third, it must be noted that the present results suggest the effect of rhythmic working memory distraction to be relatively small. In the main experiment, the maximum and minimum of average proportion correct as a function of distractor onset differed only by 0.052 (0.069 in the follow-up experiment). For comparison, the range of average recall accuracy as a function of number position, which is known to be a large effect in the Irrelevant-Speech Task, was 0.518. While we consider it likely that the former effect could be increased in future studies that employ task-irrelevant stimuli of higher distractibility, it must also be considered that the rhythmic modulation of vulnerability to distraction might be a small effect of limited practical significance.

## Conclusion

The primary goal the present study was to enhance our understanding of how the human neuro-cognitive system orchestrates the suppression of task-irrelevant distraction dynamically in time. Convergingly, the results of two experiments indicate that during retention of target information in working memory, the suppression of distractors fluctuates at a rate of ~2.5 Hz. Neurally, this effect was in part mediated by the strength of distractor encoding, where stronger encoding occurred in sync with behavioural indices of stronger distraction.

The present study sets the stage for future work to test (i) whether the observed rhythmic distractor suppression can be entrained by external stimulation; (ii) whether top-down executive control has the potency to modulate rhythmic distractor suppression to optimise task goals; and (iii) to what degree rhythmic distractor suppression is independent of rhythmic target sampling and how these two interact.

## Acknowledgements

This work was supported by the DFG (Deutsche Forschungsgemeinschaft to M.W.; grant number: WO 2371/1-1) and the European Research Council (ERC Consolidator grant to J.O.; grant number: ERC-CoG-2014-646696). The authors declare no conflict of interest. We thank Molly Henry for fruitful discussions about the analysis pipeline used in the present study. Two anonymous reviewers provided valuable feedback on an earlier manuscript version.

